# Reconstructing lost BOLD signal in individual participants using deep machine learning

**DOI:** 10.1101/808089

**Authors:** Yuxiang Yan, Louisa Dahmani, Lunhao Shen, Xiaolong Peng, Danhong Wang, Jianxun Ren, Changgeng He, Changqing Jiang, Chen Gong, Ye Tian, Jianguo Zhang, Yi Guo, Yuanxiang Lin, Meiyun Wang, Luming Li, Bo Hong, Hesheng Liu

**Affiliations:** Athinoula A. Martinos Center for Biomedical Imaging, Department of Radiology, Massachusetts General Hospital, Harvard Medical School, Charlestown, MA, USA; Department of Biomedical Engineering, Tsinghua University, Beijing, China; Department of Radiology, Henan Provincial People’s Hospital, Zhengzhou, China; National Engineering Laboratory for Neuromodulation, School of Aerospace Engineering, Tsinghua University, Beijing, China; Department of Neurosurgery, Tiantan Hospital, Capital Medical University, Beijing, China; Department of Neurosurgery, Peking Union Medical College Hospital, Beijing, China; Department of Neurosurgery, First Affiliated Hospital of Fujian Medical University, Fuzhou, China; Beijing Institute for Brain Disorders, Capital Medical University, Beijing, China; Department of Neuroscience, Medical University of South Carolina, Charleston, SC, USA

**Keywords:** machine learning, functional connectivity, fMRI

## Abstract

The blood oxygen level-dependent (BOLD) signal in functional neuroimaging suffers from magnetic susceptibility artifacts and interference from metal implants. The resulting signal loss hampers functional neuroimaging studies and can lead to misinterpretation of findings. Here, we reconstructed compromised BOLD signal using deep machine learning. We trained a deep learning model to learn principles governing BOLD activity in one dataset and reconstructed artificially-compromised regions in another dataset, frame by frame. Strikingly, BOLD time series extracted from reconstructed frames were correlated with the original time series, even though the frames did not independently carry information about BOLD fluctuations through time. Moreover, reconstructed functional connectivity (FC) maps exhibited good correspondence with the original FC maps, indicating that the deep learning model recovered functional relationships among brain regions. We replicated this result in patients whose scans suffered signal loss due to intracortical electrodes. Critically, the reconstructions captured individual-specific information rather than group information learned during training. Deep machine learning thus presents a unique opportunity to reconstruct compromised BOLD signal while capturing features of an individual’s own functional brain organization.

## Introduction

The blood oxygen level-dependent (BOLD) signal, acquired during functional magnetic resonance imaging (fMRI), is subject to a number of artifacts, such as magnetic susceptibility artifacts and interference from metal implants. For example, intracortical electrodes implanted in patients interfere with the BOLD signal, potentially due to their lead connectors, resulting in significant signal loss in brain regions close to the connection site on the skull^1^. This hampers studies that investigate whole-brain activity and functional connectivity, and may result in misinterpretation of findings. To date, there are no post-processing MRI methods that can mitigate such interference.

A newly-proposed deep machine learning model, called deep convolutional generative adversarial networks (DCGAN), provides a possible solution for reconstructing lost information^2-6^. In the DCGAN approach, two networks—a generator and a discriminator—are pitted one against the other and are trained and optimized simultaneously. Remarkably, it does not simply assemble pieces of images it was trained on, but rather generates new images that are internally cohesive. For example, DCGAN models can successfully fill in missing portions of photographs of human faces and create pictures of human faces, birds, and even art. Like photographs, BOLD images carry internally cohesive information. Embedded within resting-state data, for example, is information about BOLD signal fluctuations in each cortical surface vertex^7^, from which we can extract meaningful information such as functional connectivity and task-evoked brain activity^8^.

Here, we show that DCGAN can be harnessed to learn individual patterns of brain activity and generate BOLD signals in artificially and non-artificially compromised cortical regions. We trained a deep learning model on a dataset containing intact BOLD frames from a sample of healthy young adult participants (Fig. 1A). We used the trained model to reconstruct BOLD images, frame by frame, in an independent test dataset in which we artificially removed cortical surface regions of different sizes (Fig. 1B). Although the individual input frames did not carry information about the evolution of the BOLD signal through time, we set out on the ambitious goal of investigating the times series and functional connectivity (FC) maps extracted from the reconstructed frames. We hypothesized that the reconstructed times series and FC maps would bear high similarity to the original ones. Additionally, the large amount of resting-state data that was available enabled us to calculate individual-level functional connectivity^9,10^. We thus tested whether machine learning can be used to reconstruct individual-specific information, or whether its ability is limited to generating images based on group-level information. Finally, we tested the DCGAN model in a clinical application, where we acquired a unique dataset by collecting extensive resting-state fMRI data both before and after electrode implantation surgery in patients with Parkinson’s disease. We sought to reconstruct regions in the post-operative scans that suffered substantial interference from the deep-brain stimulation (DBS) electrodes and connectors. The availability of pre-implantation scans meant we had a reference against which to compare the reconstructed images, assuming functional connectivity is stable and unchanged by the implantation surgery. We hypothesized that the reconstructed BOLD signals from the post-surgical data would be highly similar to the pre-surgical BOLD signals, and that they would reflect patterns of activity that are specific to the individual.

**Fig. 1.**
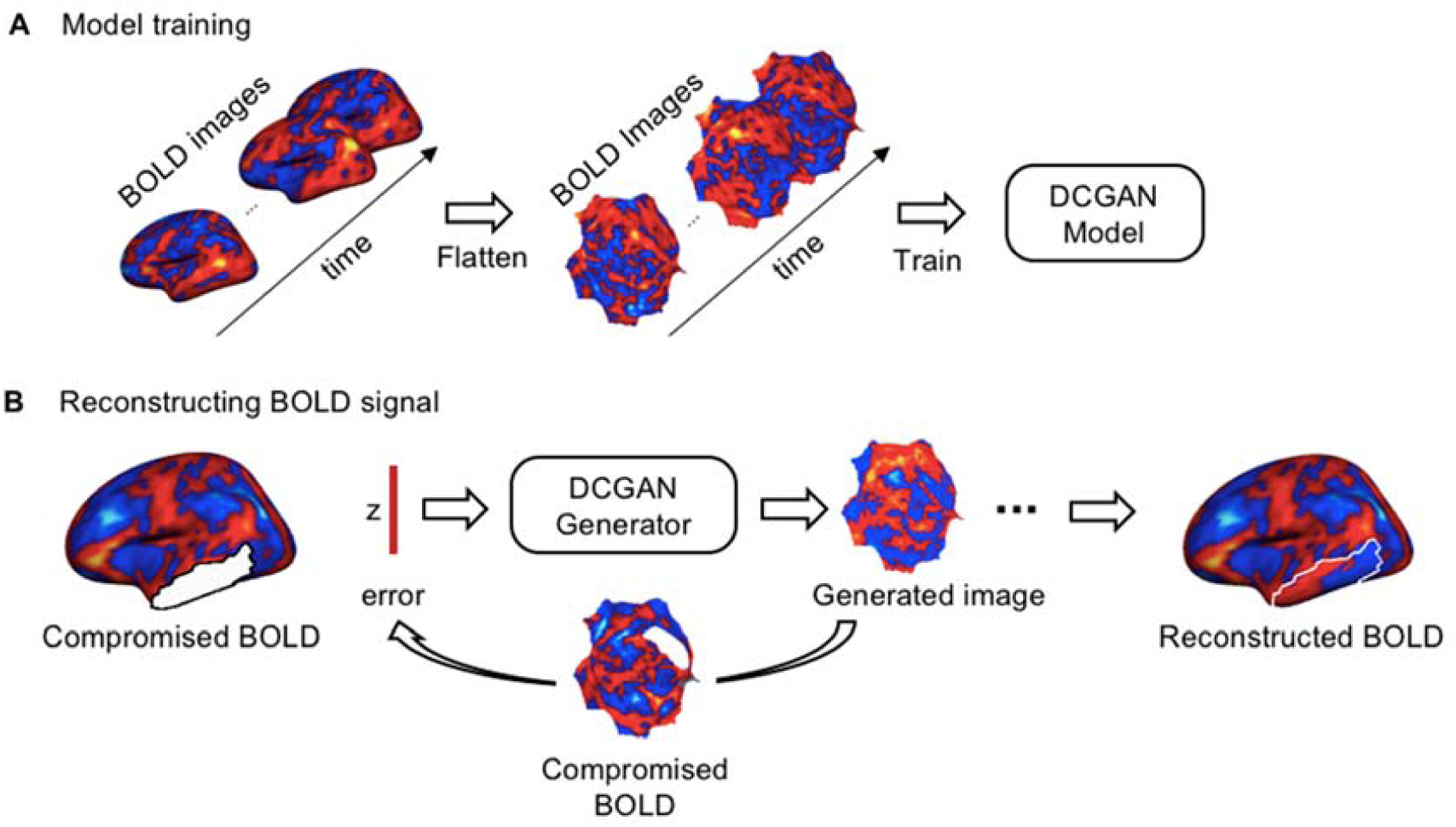
Method for reconstructing lost signal in BOLD images using deep convolutional generative adversarial networks (DCGAN). (A) 19,200 BOLD frames from 80 participants are flattened and fed to the DCGAN model for training. The DCGAN model consists of two networks, a generator and a discriminator. The generator aims to learn the distribution of the BOLD activity within the frames used as input and makes a projection from a random vector *z*, sampled from latent space *Z*, to a flattened BOLD frame *G(z)*. The discriminator is trained to distinguish the real BOLD frames from the BOLD patterns simulated by the generator. The generator and discriminator are optimized simultaneously through the two-player minimax game. (B) The trained generator is used to reconstruct the compromised BOLD frames. We first created compromised BOLD frames by removing the BOLD signal in predefined regions (here, in the temporal cortex, shown as white mask). Each compromised BOLD frame is flattened and inputted to the trained generator. Vector *z* is iteratively optimized by the gradient-descent method in the latent space to minimize the difference between the generated BOLD frames *G(z)* and the authentic data *x* outside the masked region. The vector is then projected to the flattened map *G(z*_*n*_) via the generator. Then, *G(z*_*n*_) is projected back into a 2562-vertex mesh representing the reconstructed BOLD activity in a single frame.

## Results

### Reconstructed BOLD signals are correlated with original ones

The DCGAN model was trained on a resting-state fMRI dataset of 80 randomly-chosen participants (240 frames for each participant) from the publicly-available Brain Genomics Superstruct Project (GSP) database^11^. In an independent test dataset comprised of 20 participants, we artificially removed BOLD signal in various cortical surface regions and used the trained generator to reconstruct compromised BOLD frames.

We found that the reconstructed BOLD frames appear similar to the original intact images (see Fig. 2A for an example). To quantitatively evaluate the reconstructive accuracy of the DCGAN model, we concatenated the reconstructed frames and compared the reconstructed and original time series. This is a particularly challenging endeavor, as images are reconstructed at each time point, and each frame does not independently hold temporal information. To conduct this comparison, we assessed the correlation between the original and reconstructed BOLD time series for each vertex within each of the artificially-compromised regions, located in different lobes (see *Methods* for more details), and calculated the overall average of these correlations across all participants. Strikingly, we found significant positive correlations, using multilevel linear models corrected for multiple comparisons using the Bonferroni correction: temporal cortex region: *r*=0.33±0.02 (*t*(19)=65.49, bootstrapped *p*<0.001), occipital cortex region: *r*=0.37±0.06 (*t*(19)=26.65, *p*<0.001), lateral frontal cortex region: *r*=0.16±0.07 (*t*(19)=9.74, *p*<0.001), medial frontal cortex region: *r*=0.23±0.04 (*t*(19)=23.24, *p*<0.001), and parietal cortex region: *r*=0.25±0.03 (*t*(19)=37.03, *p*<0.001). Although these correlations are low to moderate, the sheer fact that the DCGAN model was able to learn individual-specific features and capture part of the BOLD fluctuations in regions with complete signal loss is striking. Fig. 2B shows an example of a reconstruction of the lateral temporal cortex in one participant, where the correlations between the reconstructed and original time series of two randomly-selected vertices are *r*=0.67, *p*<0.001 for Vertex 1 and *r*=0.46, *p*<0.001 for Vertex 2. Importantly, within the compromised region, the reconstructed BOLD signals exhibit various patterns of activity that are not necessarily correlated with each other. For example, the correlation between the time series of the two vertices above is *r*=-0.08, *p*=0.22 (Fig. 2B).

**Fig. 2.**
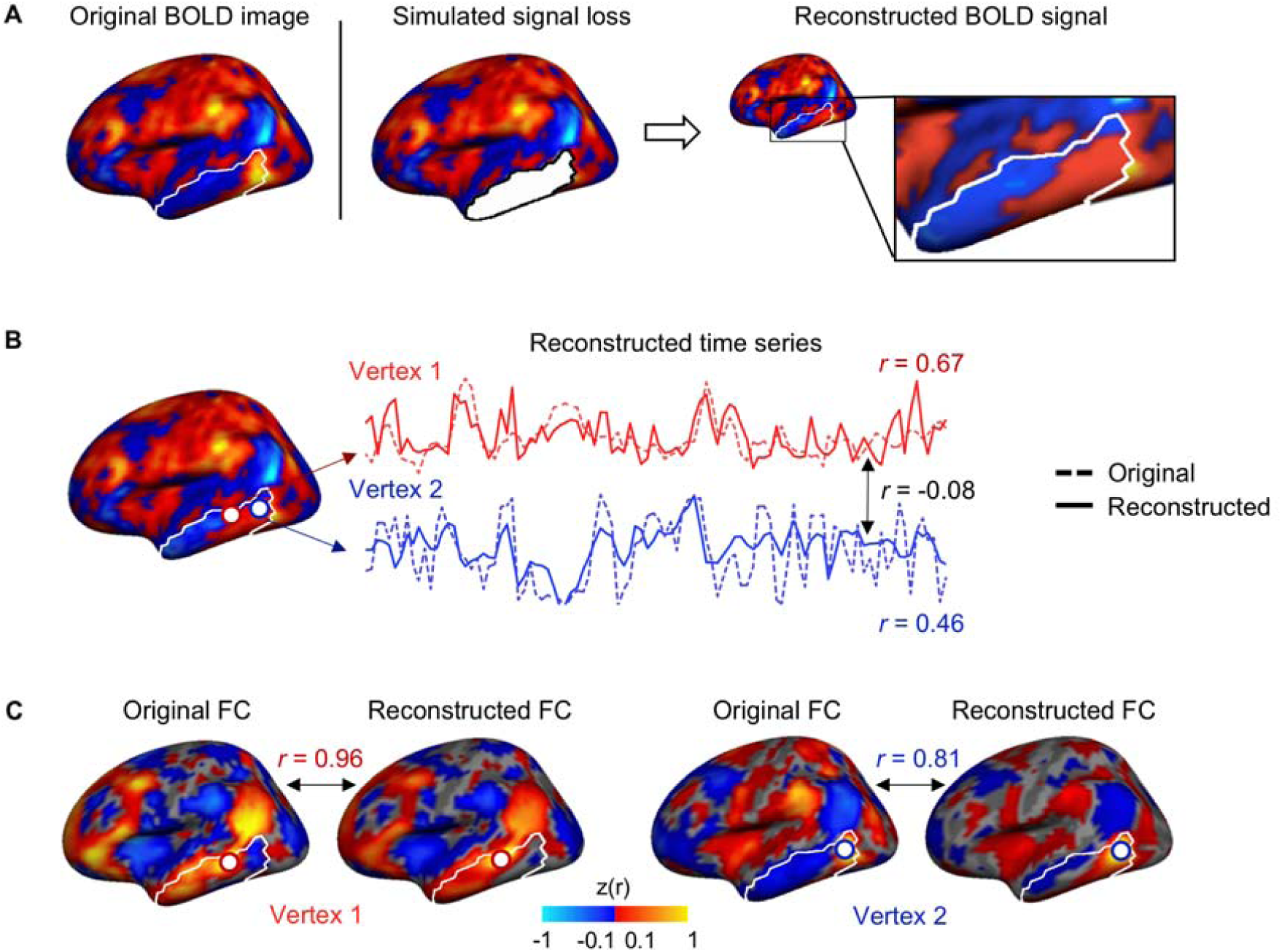
The reconstructed BOLD signals are highly similar to the original signals. (A) Here we show an example of an intact BOLD frame from a representative participant in the GSP dataset, which we used to create an artificially-compromised BOLD frame by removing the signal in a predefined region (here, a region within the temporal cortex). The compromised BOLD frame is fed to the trained DCGAN model, which then generates a reconstructed BOLD frame based on the information within the compromised frame. (B) The time series of two vertices in the reconstructed region are shown (solid lines), along with these vertices’ original time series taken from the intact BOLD frame (dashed lines). Each white circle represents a vertex. The reconstructed time series are highly similar to the original time series, as they exhibit high correlations (r = 0.67 for the left-most vertex outlined in red and r = 0.46 for the right-most vertex outlined in blue). To illustrate that the DCGAN model does not simply generate time series that follow the same fluctuations over time, we correlated the time series of the two generated vertices. The resulting correlation is r = −0.08, indicating that the DCGAN model takes into account the variability in activity between different vertices. (C) We investigated the functional connectivity (FC) maps of the original and reconstructed vertices. The two reconstructed FC maps show high similarity to the original FC maps (r = 0.96 for the left-most vertex outlined in red and r = 0.81 for the right-most vertex outlined in blue). These findings indicate that the DCGAN model is able to learn time-varying and functional connectivity characteristics of BOLD activity within individuals and to generate images that are highly realistic.

We also assessed BOLD reconstructive accuracy by generating FC maps for all reconstructed vertices and comparing them to the FC maps of the corresponding vertices in the original intact BOLD frames. As an example, in Fig. 2C, we show the similarity between reconstructed and original FC maps for the same two temporal vertices as in Fig. 2B, in a representative participant. The FC map similarity is very high, with *r*=0.96, *p*<0.001 for Vertex 1 and *r*=0.81, *p*<0.001 for Vertex 2. In Fig. S1, we show original and reconstructed FC maps sourced from one randomly selected vertex within each of the other artificially-compromised regions in a representative participant. All reconstructed FC maps yielded high similarity to the original ones: *r*=0.84, *p*<0.001 for the lateral frontal vertex, *r*=0.89, *p*<0.001 for the medial frontal vertex, *r*=0.84, *p*<0.001 for the lateral parietal vertex, and *r*=0.88, *p*<0.001 for the occipital vertex. We then evaluated the average reconstruction accuracy across all participants. When taking into account all the vertices within the compromised regions, multilevel linear models, corrected for multiple comparisons using the Bonferroni correction, revealed that the cortical FC maps of the reconstructed vertices are highly similar to those of the original vertices in all regions. The average correlation between corresponding FC maps is *r*=0.62±0.06 (*t*(19)=47.88, *p*<0.001) for the temporal cortex, *r*=0.61±0.05 (*t*(19)=51.87, *p*<0.001) for the medial frontal cortex, *r*=0.60±0.04 (*t*(19)=65.04, *p*<0.001) for the lateral frontal cortex, *r*=0.70±0.03 (*t*(19)=110.62, *p*<0.001) for the parietal cortex, and *r*=0.79±0.05 (*t*(19)=68.39, *p*<0.001) for the occipital cortex.

We also evaluated reconstructive accuracy according to the size of the compromised regions (Fig. S2A) by correlating the reconstructed and original BOLD times series. When there are no compromised regions (0% Fig. S2B), the reconstructive accuracy is very high, *r*=0.85±0.00, *p*<0.001. Once the compromised region covers 10% of the cortical surface, the mean reconstructive accuracy drops to *r*=0.51±0.07, *p*<0.001. From there, there is a steady decrease in reconstructive accuracy as the mask of compromised regions increases in size (*F*(2.62,23.55)=93.68, *p*<0.001) (Fig. S2B). When the mask size reaches 40% of the cortical surface, the reconstructive accuracy is *r*=0.34±0.09. It should be noted that a complete signal loss in 40% of the cortical surface may represent an extreme case; nevertheless, the reconstruction still recovers some important characteristics of a given individual’s functional connectivity.

### Reconstructed BOLD signals are individual-specific

We investigated whether the reconstructed BOLD signals reflect general trends in BOLD activity or whether they capture patterns of activity that are specific to the individual participant. To test this, for each vertex within a compromised region, we calculated the correlation between each individual’s reconstructed FC map and i) their original intact FC map; ii) the group-averaged FC map from the training dataset; and iii) the FC map of each participant’s most similar individual (MSI; see Methods), i.e. the individual in the training dataset that most resembles their functional connectivity patterns. In Fig. 3A, we show examples of FC maps and correlations for the same two vertices as in the BOLD time series analyses above. The reconstructed FC maps show highest correlations with the original FC maps. In Fig. 3B, we make the same comparisons but this time across all vertices within the five cortical masks combined using a repeated measures ANOVA. Reconstructed and original FC maps share many features and exhibit an average correlation of *r*=0.66±0.03, which is significantly different from 0 (*t*(19)=116.77, *p*<0.001). The average correlation between the reconstructed FC maps and the training group-averaged FC maps is *r*=0.44±0.02 and is also significantly different from 0 (*t*(19)=85.33, *p*<0.001).

**Fig. 3.**
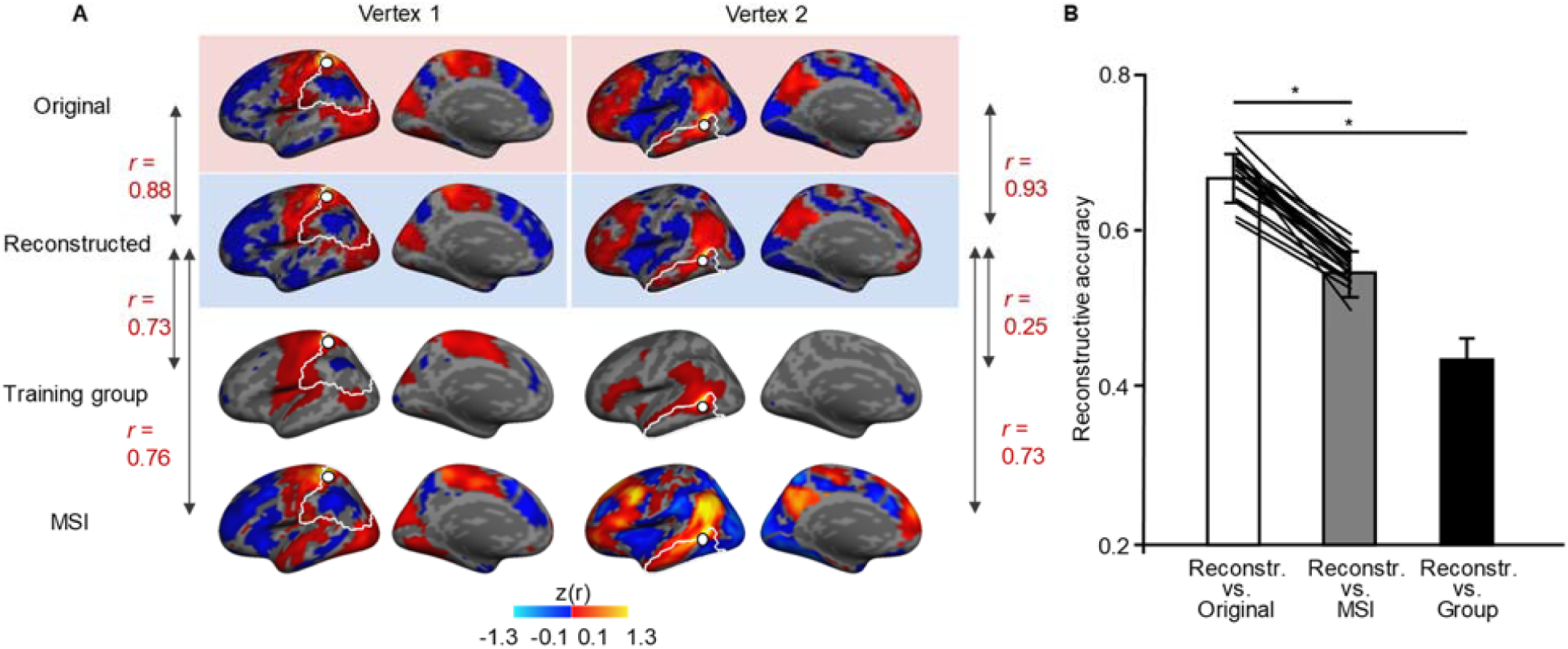
DCGAN-reconstructed functional connectivity maps capture individual-specific information. (A) For each participant in the test dataset, we generated functional connectivity (FC) maps from all vertices within each of the compromised regions. Here, we show FC maps calculated from two randomly selected seeds on the cortical surface, one in the motor region of the lateral parietal cortex and one in the lateral temporal cortex. We generated FC maps from intact BOLD frames (first row), reconstructed BOLD frames (second row), group-averaged BOLD frames (third row), and BOLD frames from each participant’s most similar individual (MSI) from the training dataset. To identify each test participant’s MSI, we correlated that participant’s FC vectors inside the compromised regions with the corresponding vectors in each of the training dataset individuals. The MSI was the individual displaying the highest average correlation. (B) If reconstructed BOLD frames capture specific details about an individual’s functional brain organization, we would expect the reconstructed FC maps to be more similar to the intact FC maps than to any other FC map. This is indeed what we found. The correlation between participants’ reconstructed and intact FC maps, averaged over all vertices within the five cortical masks (*r* = 0.66 ± 0.03), was significantly higher than the correlation between reconstructed and training group-averaged FC maps (*r* = 0.44 ± 0.02; *t*(19) = 34.53, bootstrapped *p* = 0.001) and also significantly higher than the correlation between reconstructed and MSI FC maps (*r* = 0.55 ± 0.02; *t*(19) = 20.70, bootstrapped *p* = 0.001). The diagonal lines between the first two bars join data points from the same participants. Error bars represent standard errors. * *p* < 0.001.

Finally, we compared the reconstructed FC maps of each individual in the test dataset to the FC maps of their MSI. The average correlation between reconstructed and MSI FC maps is *r*=0.55±0.02, which is significantly different from 0 (*t*(19)=116.86, *p*<0.001). We statistically compared these correlations and found them to be significantly different from one another (*F*(2,38)=927.69, *p*<0.001). Post-hoc tests (bootstrapped paired t-tests) revealed that the correlation between reconstructed and original FC maps is greater than all other correlations (reconstructed vs. training group-averaged maps: *t*(19)=34.53, bootstrapped *p*=0.001; reconstructed vs. MSI maps: *t*(19)=20.70, bootstrapped *p*=0.001). The fact that the correlation between reconstructed and original FC maps is higher than the correlation between reconstructed and training dataset FC maps indicates that during the training process, the generator did not simply learn general trends in BOLD activity but was able to infer the co-activating patterns from the individual-specific BOLD frames used as input in the test phase. Critically, the reconstructed BOLD FC maps are more representative of each individual’s own specific patterns of functional connectivity than of any other individual in the training dataset, indicating that the DCGAN model is able to capture individual-specific information about functional brain organization.

### Signals are successfully reconstructed in a clinical sample

We tested the DCGAN model in patients whose MRI signals were interfered with by metal implants. In patients with Parkinson’s disease undergoing DBS^12^, intracortical electrodes were implanted to stimulate the brain over the long term^13^. However, wires outside the skull connecting the simulator to the implanted electrodes strongly interfere with the acquisition of the BOLD signal during post-surgical fMRI studies, resulting in signal loss in temporal, parietal, and occipital regions (Fig. 4) and in abnormal measurements of functional connectivity. We used the DCGAN model to reconstruct BOLD signals in cortical regions that were compromised by the implantation. As a proof of concept, two patients with extensive resting-state fMRI data both before and after electrode implantation surgery were included in this analysis. We first identified the compromised region where vertices exhibited a sharp contrast in signal amplitude before and after the electrode implantation surgery. The compromised region covered 7.5% of the cortical surface in one patient and 7.8% in the other.

**Fig. 4.**
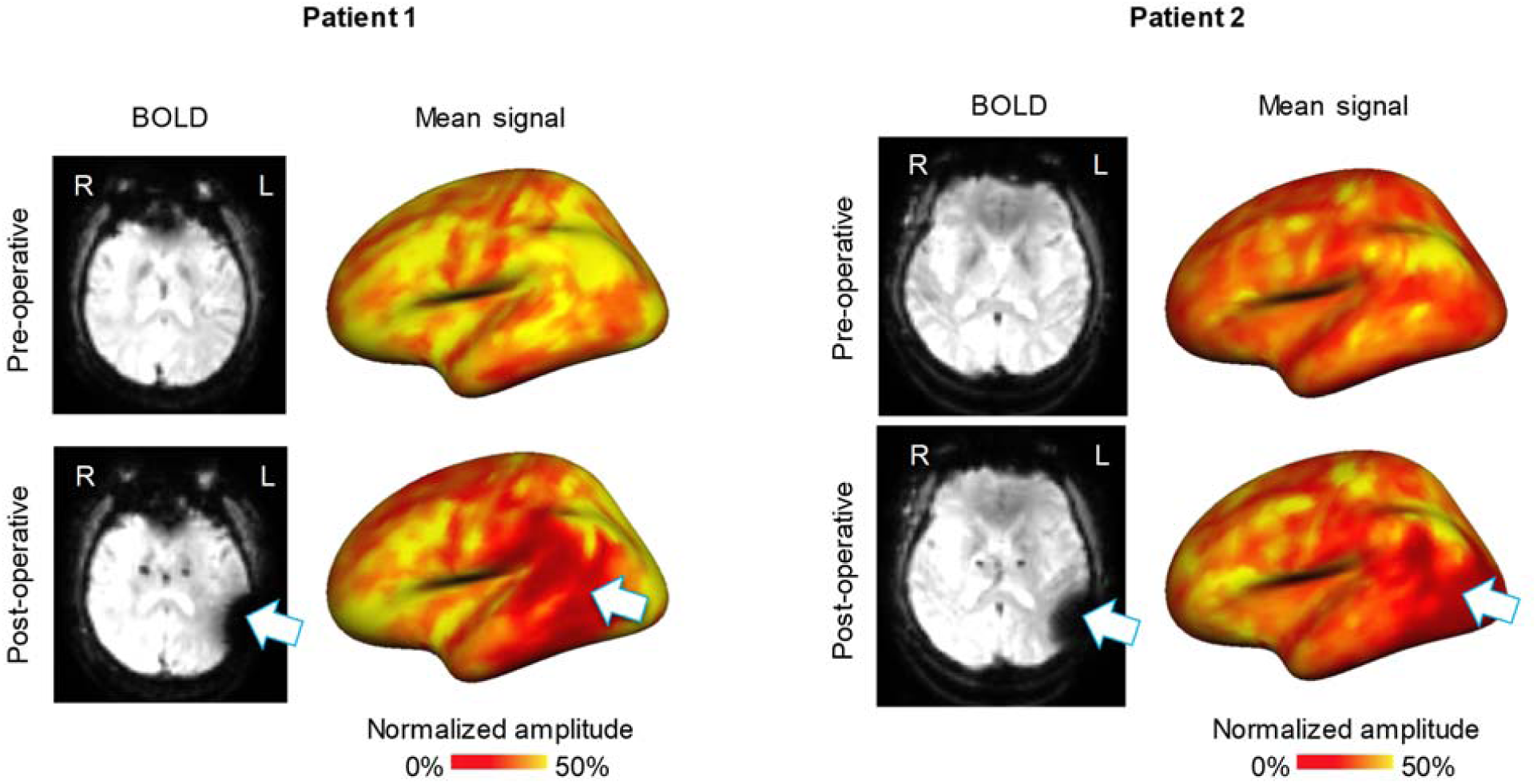
Implanted electrode interference severely reduces BOLD signal amplitude. **Pre- and post-operative BOLD frames are shown in the left panels for each patient with P**arkinson’s disease. Patients had an electrode implanted in the subthalamic nucleus, resulting in interference around left temporal, parietal, and occipital regions. The pre- and post-operative normalized and averaged BOLD signals are shown on inflated cortical surfaces in the right panels. Following surgery, the BOLD signal amplitude is severely affected in regions close to the electrode wires (temporal and parietal regions for Patient 1; temporal, parietal, and occipital regions for Patient 2).

Once the DCGAN model reconstructed the BOLD signals in the compromised regions, we investigated whether there was any residual loss within these regions. We compared the normalized amplitudes of the pre-operative BOLD signal with that of i) the post-operative compromised BOLD signal (Fig. S3A, left), and ii) the post-operative reconstructed BOLD signal (Fig. S3A, right). While the post-operative signal demonstrated substantial attenuation within the DBS-compromised cortical regions, the reconstruction displayed no residual signal loss. Using the pre-operative average signal amplitudes as reference, we calculated the normalized BOLD amplitudes (%) of the masked-out vertices in the post-operative images and reconstructed images. A multilevel linear model shows that the BOLD amplitudes are significantly different under pre-operative, post-operative, and reconstructed conditions, *F*(2,392)=4571.06, *p*<0.001. Examination of the parameter estimates revealed that the post-operative BOLD amplitudes are substantially and significantly attenuated in comparison to the pre-operative BOLD amplitudes (*t*(392)=-79.09, *p<*0.001). The reconstructed BOLD signals show negligibly higher amplitudes compared to the pre-operative BOLD signals (*t*(392)=2.31, *p*=0.02). The average pre-operative, post-operative, and reconstructed normalized BOLD amplitudes are shown in Fig. S3B.

Next, we sought to evaluate the reconstructive accuracy using functional connectivity analyses, similar to those conducted in the healthy cohorts above. We did not consider BOLD time series here as the pre- and post-operative fMRI scans were obtained at different time points. However, functional brain organization, as assessed with functional connectivity, is assumed to be relatively stable over time^14^. Although the implantation surgery may cause microlesion effects and lead to changes in brain circuits involving the stimulation target (i.e., the subthalamic nucleus), the surgery is unlikely to change functional connectivity in the area of signal loss, which is relatively far from the location of the stimulator and the motor network being modulated. To support this, we investigated functional connectivity of the right temporoparietal region of the two patients (note that signal loss was observed only in the left hemisphere). We found that the post-operative FC maps in the right hemisphere are highly and significantly correlated with the pre-operative FC maps (*r*=0.78±0.01; *t*(1)=70.45, *p*=0.009). However, for the compromised region in the left hemisphere, we found that the post-operative FC maps are also positively correlated with the pre-operative FC maps, but this correlation is not significantly different from 0 (*r*=0.42±0.07; *t*(1)=5.73, *p*=0.11). As an example, we show cortical FC maps using a seed placed in each patient’s compromised region (Fig. S4). Unlike the weak and disorganized FC maps obtained from patients’ compromised left temporoparietal region, the FC maps generated from seeds in the right temporoparietal region show high similarity to their pre-operative FC maps (right hemisphere seeds across both patients: *r*=0.83±0.03, *p*<0.001; left hemisphere seeds across both patients: *r*=0.38±0.13, *p*<0.001) (Fig. S4).

Having shown that FC maps are relatively stable following electrode implantation, we next assessed the reconstructive accuracy of our DCGAN model for the patients’ compromised region in the left hemisphere. A multilevel linear model that took into account all the vertices inside the compromised regions showed that the reconstructed post-operative FC maps are positively correlated with the pre-operative FC maps (*r*=0.54±0.02), a correlation significantly different from 0 (*t*(1)=28.72, *p*=0.022). As mentioned above, the post-operative FC maps were also positively correlated with the pre-operative FC maps, but this correlation was not significantly different from 0 (*r*=0.42±0.07; *t*(1)=5.73, *p*=0.11). Another multilevel linear model showed that the correlations between reconstructed and pre-operative FC maps are significantly higher than the correlations between post-operative and pre-operative FC maps (*t*(392)=-9.22, *p*<0.001). As an example, we show cortical functional connectivity maps using a seed placed in each patient’s compromised region (Fig. 5, same seeds as in Fig. S4). As expected, the FC maps obtained from patients’ compromised post-operative BOLD images did not capture the connectivity patterns observed in the pre-surgical data. However, the FC maps generated from the reconstructed BOLD images show high similarity to the pre-operative FC maps (reconstructed vs. pre-operative across both patients: *r*=0.61±0.01; post-operative vs. pre-operative across both patients: *r*=0.38±0.13). These results indicate that the BOLD signals reconstructed in the compromised regions are representative of the patients’ intact functional connectivity patterns.

**Fig. 5.**
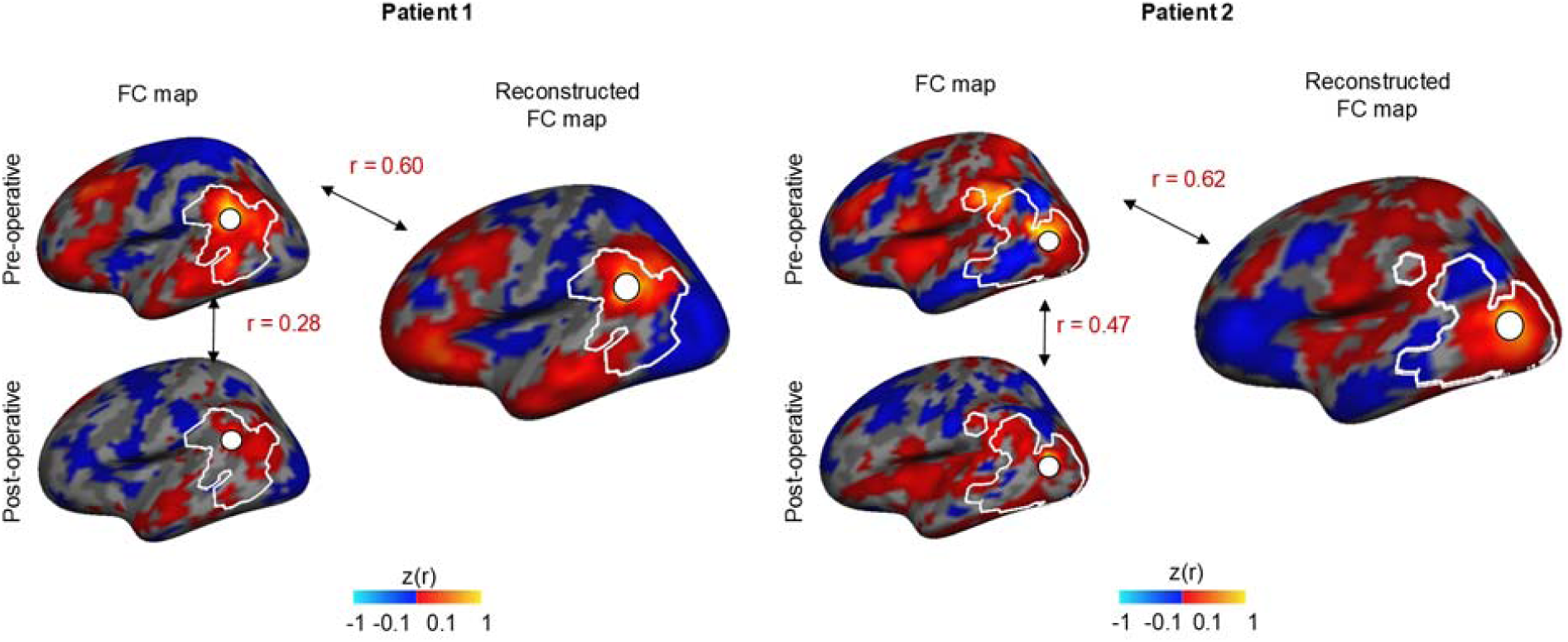
The DCGAN model can reconstruct BOLD signals compromised by intracortical electrode interference in patients with Parkinson’s disease. We generated the FC map of a given vertex before and after surgery in each patient (left panels). Delineated in white is the region that is affected by the electrodes, defined by identifying the vertices which showed a stark decrease in BOLD amplitude after the implantation surgery. The post-operative FC maps are substantially different from the pre-operative FC maps due to the intracortical electrodes, with a relatively low correlation of *r* = 0.28 in Patient 1 and *r* = 0.47 in Patient 2. We also generated FC maps for the same vertices, this time using the reconstructed BOLD frames (right panels). The generated FC maps are highly similar to the intact pre-operative FC maps, as indicated by a correlation of *r* = 0.60 for Patient 1 and *r* = 0.62 for Patient 2. Thus, the DCGAN model is able to reconstruct patients’ functional brain organizations.

## Discussion

The current study aimed to reconstruct fMRI BOLD signal inside cortical regions that suffered signal loss due to various artifacts. We used deep convoluted generative adversarial networks (DCGAN), a recent advance in machine learning algorithms, to leverage functional information embedded within BOLD frames and reconstruct the signal in compromised regions, frame by frame. We reconstructed BOLD signals in two cohorts: healthy young adults with artificially-compromised cortical regions and patients whose fMRI scans suffered from interference due to metal implants. Our results indicate that such a machine learning technique successfully reconstructs individual-specific BOLD signals and can approximate the functional connectivity patterns observed in the intact or unimpaired state.

The missing BOLD signal was reconstructed frame by frame, following which we modeled the time series for all individual vertices whose signal was compromised. We found the reconstructed time series to be similar to the original time series. This indicates that our model was able to recover dynamic brain activity over time, at the level of single vertices within individual participants. The same was found for the reconstructed and original functional connectivity maps. These findings are striking, as the images were reconstructed at each time point independently and single input frames did not carry time-varying information. The generator was thus able to learn information beyond what was presented at face value during the test phase, and modeled accurate functional interactions between brain regions and the changing dynamics of the BOLD signal through time.

The reconstruction of the compromised signals is based on learning the functional relationships between different brain regions in intact BOLD frames from a large dataset. The trained model deduces possible signal distributions in the compromised region using the remaining intact BOLD amplitude patterns in the individual. Thus, the generator learns functional activity patterns that are specific to each individual and builds a high-dimensional space that is sensitive to variations across individuals. Indeed, we show that the reconstructed BOLD signals capture individual differences in patterns of activity. A given individual’s reconstructed FC maps were more similar to their original FC maps than to the training group-averaged maps or to the ones belonging to the most similar individual from the training dataset. Therefore, the deep machine learning model may be useful in recovering lost signal in clinical settings, where the focus is on the individual.

On that note, our method has several potential clinical applications. Revealing individual-based functional activity is critical not only for understanding the functional network organization of the human brain^15^, but also for personalized medicine, such as when precise cortical mapping is required for neurosurgery or neuromodulation^16,17^. Our individual-specific machine learning method can, as we have shown, reconstruct BOLD signal that was lost due to intracortical electrode interference. We showed that the model generates FC maps with high reconstructive accuracy, as they exhibit high similarity to maps derived from presurgical images. Thus, machine learning-based reconstruction can impact the investigation of various disorders where intracortical electrodes are used for diagnosis or therapy, such as in epilepsy, Parkinson’s disease, depression, obsessive compulsive disorder, among others. This method could also help mitigate registration problems in fMRI that occur with data from patients with brain lesions due to stroke or tumor resections, by filling in the compromised regions prior to registration. It could also fill in regions that are routinely cut off during acquisition, such as the top of the brain or the lower portion of the cerebellum. Finally, in both clinical and non-clinical settings, machine learning could be used to reconstruct the BOLD signal in regions that are susceptible to signal loss and geometric distortion, such as the orbitofrontal cortex and temporal cortex. However, this could only be done once the model can be trained on uncompromised images, thus appropriate data acquisition methods that can counteract these susceptibility artifacts will first have to be developed.

We observed that the reconstructive accuracy differed from region to region. Reconstructive accuracy may be affected by a number of factors. One factor is the size of the compromised region, as smaller regions yield better reconstructions. A second factor is the shape of the region, as closer proximity to uncompromised vertices is likely to result in better reconstruction. A third factor is whether the affected region has important large-scale network connectivity, which would increase accuracy as fMRI activity outside of the compromised region would bear information relevant to the activity within the compromised region. Additionally, the distribution of learned functional activation patterns is constrained by the training dataset. More efforts are necessary to evaluate the reconstructive accuracy of the model when the compromised BOLDs are not acquired using the same MRI and scanning parameters as the training data. Another limitation of our study is that the DCGAN model dealt with 2D images. Currently, it is not able to reconstruct whole-brain 3D images as the computational power required is too high. We hope that technological advances will soon enable this type of modelling. In the meantime, it may be possible to reconstruct signal in small 3D volumes.

Besides 3D modelling, we propose one area of future study to improve the current machine learning model, which would be to consider the causal interactions across time frames. Currently, each of the BOLD frames is used separately to supervise the learning process of the adversarial networks. In this way, the connections among cortical areas, which are not included in single BOLD frames, are difficult to detect. Feeding DCGAN a combination of BOLD frames may improve their modelling power.

In summary, we harnessed the learning power of deep convolutional neural networks to generate BOLD signals in regions that experienced signal loss. We have shown that it is possible to reconstruct lost BOLD signal in a healthy population as well as in a clinical sample with compromised fMRI. In both cases, the reconstructed signals closely resemble the uncompromised signals. Notably, the reconstructed signals are coherent with each individual’s functional brain organization. Such a method could benefit personalized clinical and non-clinical studies where brain regions suffer signal dropout, distortion, or deformation.

## Materials and Methods

### Participants

We used the resting-state fMRI data of 100 healthy young adult participants from the GSP dataset (mean age: 22.0 ± 3.2 years; 50 women, 50 men)^11^. We also recruited two patients (two men, aged 46 and 51) who had intracortical electrodes implanted for deep-brain stimulation (DBS) treatment of Parkinson’s disease. The patients underwent stereotactic implantation of quadripolar DBS electrodes (PINS Medical, Model L301C) in the subthalamic nucleus. Microelectrode recording and stimulation guided the electrode implantation, and the electrodes were connected to extension leads (PINS Medical, Model E202C), which were themselves connected to the implanted pulse generator (PINS Medical, Model G106R). All participants provided written informed consent in accordance with guidelines set by the Institutional Review Boards of Harvard University, Partners Healthcare, or Beijing Tiantan Hospital of Capital Medical University.

### MRI data acquisition

Each healthy young participant from the GPS dataset underwent one structural scan and two resting-state fMRI scans (6 min and 12 s per scan). Data were collected on matched 3T Tim Trio scanners (Siemens, Erlangen, Germany) using a 12-channel phased-array head coil. Structural data included a high-resolution multi-echo T1-weighted magnetization-prepared gradient-echo image (TR = 2,200 ms, TE = 1.54 ms for image 1 to 7.01 ms for image 4, TI = 1,100 ms, flip angle = 7°, voxel size: 1.2 × 1.2 × 1.2 mm, FOV = 230, slices = 720). Resting-state fMRI images were acquired using the gradient-echo echo-planar pulse sequence (TR = 3,000 ms, TE = 30 ms, flip angle = 85°, voxel size: 3 × 3 × 3 mm voxels, FOV = 216, slices = 47 slices collected with interleaved acquisition with no gap between slices). Whole-brain coverage including the entire cerebellum was achieved with slices aligned to the anterior commissure-posterior commissure plane using an automated alignment procedure, ensuring consistency across participants^18^. Participants were instructed to stay awake, keep their eyes open, and minimize head movement; no other task instruction was provided.

Each of the two patients with Parkinson’s disease underwent four resting-state fMRI scans (6 min 8 s per scan) at two time points: at baseline before the DBS electrode implantation surgery and one month after. The patients were instructed to stay awake and keep their eyes open. The DBS electrodes were turned off during post-surgical fMRI scanning. The specific absorption rate-estimated values were continuously monitored throughout the scanning sessions. MRI data was collected on a Philips Achieva 3.0 Tesla TX whole body MR scanner using a 32-channel receive-only head coil. Structural images were acquired using a sagittal magnetization-prepared rapid gradient echo T1-weighted sequence (TR = 7.6 ms, TE = 3.7 ms, TI = 1000 ms, flip angle = 8°, voxel size = 1 × 1 × 1mm, FOV = 256, slices = 180). Functional data was collected using an echo planar imaging sequence (TR = 2000 ms, TE = 30 ms, flip angle = 90°, voxel size=2.875 × 2.875 × 4 mm, FOV = 230, slices = 37).

### Data processing

Structural data were processed using FreeSurfer version 4.5.0. Surface mesh representations of the cortex from each individual participant’s structural images were reconstructed and registered to a common spherical coordinate system^19^. The structural and functional images were aligned using boundary-based registration using the FsFast software package (http://surfer.nmr.mgh.harvard.edu/fswiki/FsFast)^20^. The preprocessed resting-state fMRI data were then aligned to the common spherical coordinate system via sampling from the middle of the cortical ribbon in a single interpolation step^21^. We registered each individual’s fMRI data to the FreeSurfer template which consisted of 40,962 vertices in each hemisphere. A 6-mm full-width half-maximum (FWHM) smoothing kernel was applied to the fMRI data in the surface space. The smoothed data were downsampled to a mesh of 2,562 vertices in each hemisphere using the mri_surf2surf function in FreeSurfer.

Resting-state fMRI data were processed using the procedures described in previous work^21-23^. The steps were as follows: i) slice timing correction (SPM2; Wellcome Department of Cognitive Neurology, London, UK); ii) rigid body correction for head motion with the FSL package^24,25^; iii) normalization of global mean signal intensity across runs; and iv) bandpass temporal filtering (0.01 – 0.08 Hz), head-motion regression, whole-brain signal regression, and ventricular and white-matter signal regression in a single step. After preprocessing, each participant’s resting-state fMRI data were normalized to [-1,1] by dividing the BOLD amplitude of each vertex by the maximum absolute BOLD value observed in each session. The normalized 2562-vertex mesh of the BOLD frames were then flattened to 2-dimensional maps using the tksurfer and mris_flatten functions in FreeSurfer^19^.

### Machine learning

We used DCGAN to reconstruct lost or compromised BOLD information. The neural network modeling was conducted in three steps. In the first step, the training phase, two competing models are trained: a generator and a discriminator (Fig. 1B). The generator is trained to encode BOLD information by feeding it intact BOLD frames from the training dataset. Using information within these frames, the generator creates new frames, which the discriminator then classifies as being either authentic (real BOLD frame) or artificial (generator-created BOLD frame). Both the generator and discriminator simultaneously continue training with new frames, and through many iterations each becomes optimized. Training ends when an optimized discriminator classifies the generated frames into one category or the other at chance level. In the second step, we created artificially-compromised BOLD frames by removing the BOLD signal within certain predefined regions, using data from the test dataset. In the third step, the signal in the compromised region is reconstructed by feeding the compromised BOLD images to the DCGAN generator (Fig. 1C). The generator then produces new complete frames based on these. Using the mask of the compromised region, the region with reconstructed signal from the newly generated frame replaces the one in the compromised frame to form a complete BOLD frame which includes the original information (BOLD signal outside of the compromised region) and the newly generated information (BOLD signal inside the reconstructed region). Following this, we evaluated the similarity between the reconstructed BOLD information and the original intact BOLD information. Each of these steps is described in more detail below.

#### Step 1: Training a generative network to encode BOLD information

Eighty participants were randomly selected from the Brain GSP dataset^11^ to build the training dataset, which consisted of 19,200 intact flattened BOLD frames. The data from 20 other participants, again selected randomly from the GSP, constituted the test dataset, which was independent from the training dataset. We used a DCGAN model^2^ to create BOLD frames based on each individual participant’s data. The generator’s goal is to create images similar enough to the original images that the discriminator is forced to randomly classify them as authentic or simulated, while the goal of the discriminator is to correctly classify images as either authentic or simulated.

In mathematical terms, the generator (*G)* samples data *x* from the true data population *p*_*data*_ and produces parameters. It maps these parameters onto a random vector *z*, which is sampled from latent space *Z*, and creates artificial images *G(z)*, which are part of the generated distribution *p*_*g*_. When the discriminator (*D)* detects a difference between the distributions *p*_*data*_ and *p*_*g*_, the generator *G* tweaks its parameters and generates images that are more similar to the authentic images. This process is repeated until the generator produces a generated distribution *p*_*g*_ that so closely matches the true data distribution *p*_*data*_ that the discriminator *D* is unable to detect a difference and classifies the authentic or generated images *G(z)* randomly.

Convolutional neural networks (CNN) were used to build the generator and discriminator^4^. During the adversarial training process, the generator and discriminator were trained simultaneously. They were optimized using a Nash equilibrium of costs two-player minimax game with value function *V(G, D)*:

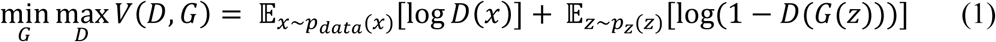

The input *z* was a sample taken from 100 dimensional uniformly distributed noise; in each dimension, the value varied from −1 to 1. The generator projected the input *z* to a small convolutional representation, and then converted the representation into a 500 × 500 pixel image through four-layer fractionally-strided convolutions^4^. The discriminator estimated the input images through four-layer-strided convolutions, and fed the layers into single sigmoid outputs. Rectified Linear Unit activation was used in the generator’s layers, except for the output layer, which used the Tanh function^26^. Leaky rectified activation was used in all of the discriminator’s layers^27^. A 64-size batch normalization was used in the training procedure for stabilization^28^, and the sigmoid cross entropy was calculated to measure the probability difference between two images. We used an Adam optimizer during the optimizing procedure of the generator and discriminator^29^, with a learning rate of 0.0002. The generator’s parameters were adjusted twice during each iteration to balance the learning speed between the generator and the discriminator.

#### Step 2: Creating the compromised BOLD frames

To allow the DCGAN model to generate BOLD signals in compromised regions, we created frames in which we removed the BOLD signal in predefined regions. We aimed to test the generator on five regions spread across the cortical surface: the lateral frontal cortex, the medial frontal, cortex, the lateral parietal cortex, the lateral temporal cortex, and the occipital cortex. These were delineated using FreeSurfer’s Desikan-Killiany atlas^30^. The Desikan-Killiany atlas labels of the artificially-compromised regions are: 4, 13, 19, 20, 21, 28 for the lateral frontal cortex (329 eliminated vertices, 12.8% of cortical surface); 3,15, 27, 29 for the medial frontal cortex (223 eliminated vertices, 8.7% of cortical surface); 9, 30, 32 for the lateral parietal cortex (429 eliminated vertices, 16.7% of cortical surface); 10 and 16 for the lateral temporal cortex (140 eliminated vertices, 5.5% of cortical surface); and 6, 12, 14, 22 for the occipital cortex (188 eliminated vertices, 7.3% of cortical surface). Masks *M* were created separately to mask out each of the eliminated regions and leave intact the other parts of the flattened BOLD activation map. After preprocessing, BOLD signals within the masks were artificially set to 0 to create the compromised BOLD frames.

We also sought to evaluate the reconstructive accuracy of our model according to the size of the compromised regions. To do this, we created ten sets of incrementally larger masks (Fig. S2A). We selected at random 10 vertices among all 2562 cortical surface vertices, located in various cortical regions. Each of these vertices served as the center of its mask set. Each mask set was comprised of six masks, whose coverage went from 10% to 60% of the cortical surface, with incremental steps of 10% (i.e., 10%, 20%, 30%, 40%, 50%, 60%). The BOLD signal inside these masks was eliminated to create 10 sets of BOLD images with increasingly larger compromised regions.

Finally, for the patients with Parkinson’s disease, we created masks which encompassed the regions in which interference was observed due to the implanted DBS electrodes. We first calculated the absolute values of the surface-based BOLD signal for each vertex, which were then averaged and normalized to [0,1] to achieve a normalized BOLD signal strength for each vertex before and after implantation surgery. Then, the pre- and post-operative BOLD amplitude maps were contrasted and the vertices with strongly reduced amplitude post-operatively were extracted to create a mask. The compromised regions comprised 193 vertices (7.5% of cortical surface) in one patient and 200 vertices in the other (7.8% of cortical surface).

#### Step 3: Reconstructing compromised BOLD signals

The trained generator was used to reconstruct the BOLD frames with artificially-compromised regions, by generating an image *G(z)* with maximal similarity to the original BOLD activation *x* in the cortical regions outside of the mask. The generator’s loss function serves to calculate the divergence between the real and simulated BOLD frames, and was defined as:

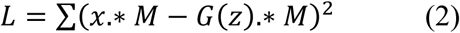

In order to minimize the loss function, an optimized *z* must be found in the latent space. After random sampling, *z* was mapped to *G(z)* by the trained generator. The location of *z* was rearranged iteratively during the optimization process using the gradient descent method to find the most similar generated image *G(z)* with minimal *L* to *x*. The number of iterations was set to 500, with a 0.000002 learning rate.

In the reconstruction step, we used 4,800 BOLD frames from the test dataset of 20 participants. The reconstructed BOLD frames were built from the masked-out flattened images. To do this, we first had to determine the spatial location of each vertex on a flattened BOLD amplitude map. Each vertex was generated and subsequently projected onto a flattened cortical surface map. The coordinates with maximum value in the flattened map were regarded as the location of the generated vertex. Using this method, we determined the correspondence between each of the vertices and their spatial location on the flattened map. This allowed us to then take the flattened map and to project it back into vertices on a BOLD frame. For each participant, the simulated BOLD frames were assembled temporally to reconstruct the BOLD time series.

### Statistical Analysis

The reconstructive accuracy of the DCGAN model was evaluated by calculating correlations between original and reconstructed BOLD signal time series and functional connectivity. The similarity between the original and reconstructed BOLD time series was only investigated in the first part of the study with healthy adults. The reconstructed signal was compared to the original signal. For the patient study, time series obtained at different time points cannot be directly compared, for this reason we did not compare the post-operative reconstructed time series to the pre-operative time series in the patients with Parkinson’s disease. As the brain’s functional connectivity (FC) patterns are relatively stable through time, we compared the FC maps generated from various seeds inside the compromised regions, in both the healthy adult dataset and the Parkinson’s disease dataset. To show that functional connectivity is relatively stable and unaffected by electrode implantation surgery, we also generated FC maps using seeds in uncompromised regions.

For the time series comparisons at any given vertex, the similarity between reconstructed and original signals was quantified by calculating the Pearson’s correlation between the two time series. The correlation values within a given masked region were then averaged across participants to represent the similarity between the reconstructed and original BOLD signals in that region.

The cortical FC of the original and reconstructed BOLD signals was also compared. For the original BOLD images, a FC map was created by calculating the *z* value of the correlation between the BOLD signal at a given vertex and the BOLD signals at all the vertices in the two cerebral hemispheres. For the reconstructed BOLD images, the compromised regions were filled with the reconstructed BOLD signal to calculate the functional connectivity of the whole brain. The vectors storing the functional connectivity of the original and reconstructed BOLD signals for a given vertex were then correlated to determine their similarity. For each of the five cortical masks, the statistical significance for the time series and FC map correlations was assessed using multilevel linear models with vertices nested within participants. Whenever the assumption of normality was violated, bootstrapped *p* values were calculated. Multiples comparisons were corrected for using the Bonferroni correction.

To assess whether the reconstructed BOLD signals were representative of individuals’ own BOLD signals or whether they simply reflected general trends in BOLD activity learned from the training dataset, we calculated the correlation between the reconstructed BOLD FC map from the test dataset and the group-averaged FC map from the training dataset in corresponding vertices. The correlations were calculated using all vertices within the five cortical masks combined.

We also compared the reconstructed FC map of each participant in the test dataset to the intact FC map of the individual in the training dataset that most resembles them, called the most similar individual (MSI). To determine each participant’s MSI, we correlated a given individual’s functional connectivity vectors for the vertices inside all five cortical masks combined with the vectors of corresponding vertices in each of the 80 training dataset participants. The training dataset individual showing the highest similarity (highest correlation) to a given participant from the test dataset was identified as that participant’s MSI. The statistical significance of each of these average correlations (reconstructed vs. original, reconstructed vs. training, reconstructed vs. MSI) was assessed by examining parameter estimates generated from a repeated measures analysis of variance (ANOVA), which also served to statistically compare whether these three average correlations were different from one another. Finally, we used paired t-tests as post-hoc tests to determine which pairs of correlations demonstrated a significant difference. We also used a repeated measures ANOVA to determine whether the size of the compromised region significantly affected the reconstructive accuracy of the generated frames across 10 cortical masks.

Means are presented along with their standard deviations (mean ± s.d.) in the results section.

## Author contributions

H.L & Y.Y designed the research; Y.Y., L.D., L.S., X.P., D.W., J.R., C.H., C.J., C.G., Y.T., J.Z., Y.G., Y.L. performed the research, Y.Y., L.D., H.L. analyzed data, L.D., Y.Y., H.L., M.W., L.L., B.H. wrote and improved the paper.

## Data and materials availability

The data that support the findings of this study are available from the corresponding author upon reasonable request.

## Supplemental Figures

**Fig. S1.**
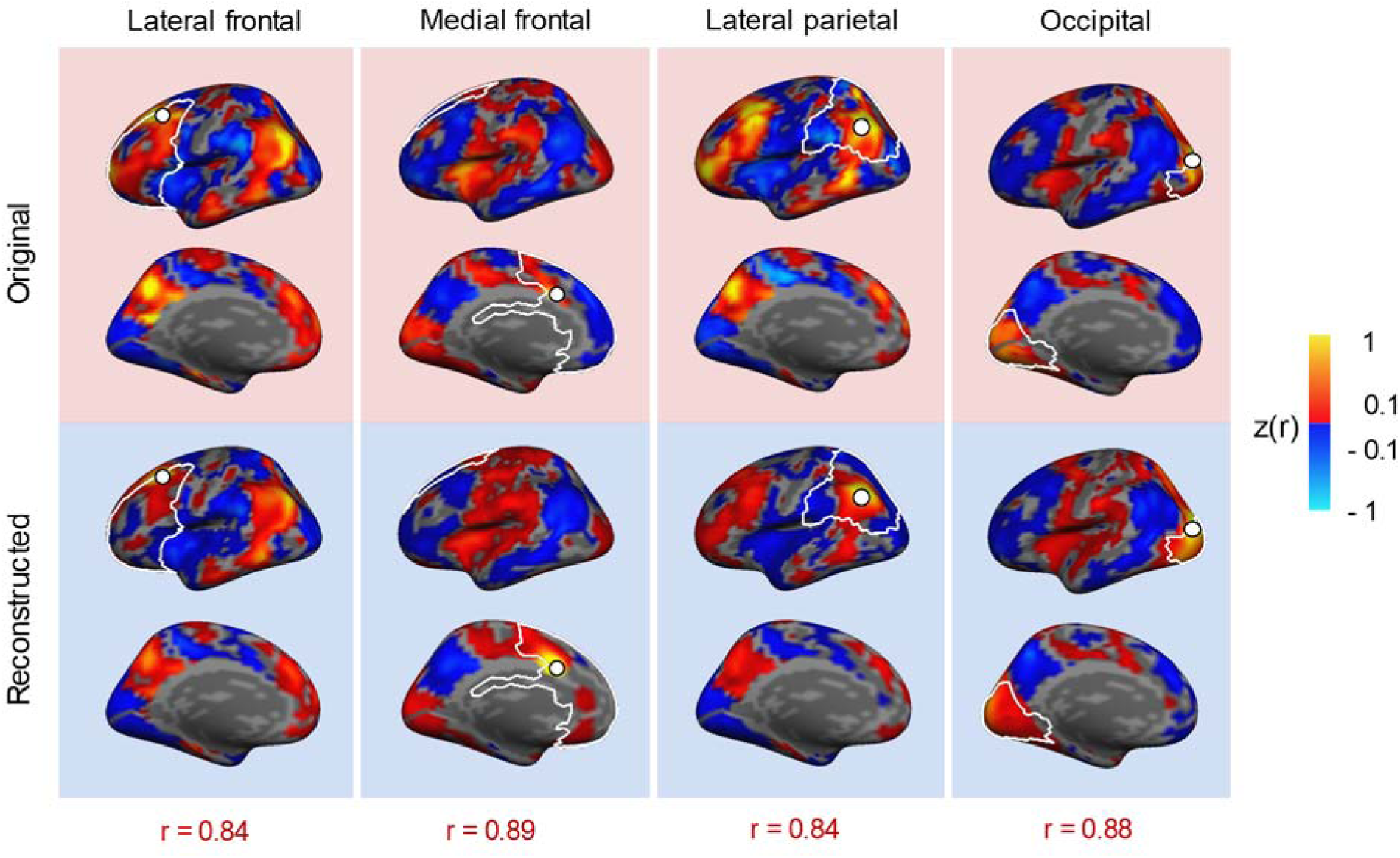
The DCGAN-generated functional connectivity maps are highly similar to the original maps, throughout cortex. The top row shows functional connectivity (FC) maps of seeds from intact BOLD frames in the lateral frontal, medial frontal, lateral parietal, and occipital cortices. The bottom row shows the FC maps of the same seeds from reconstructed BOLD frames. The similarity between the original and reconstructed FC maps is high, with the following correlation coefficients: *r* = 0.84 for the lateral frontal seed, *r* = 0.89 for the medial frontal seed, *r* = 0.84 for the lateral parietal seed, and *r* = 0.88 for the occipital seed.

**Fig. S2.**
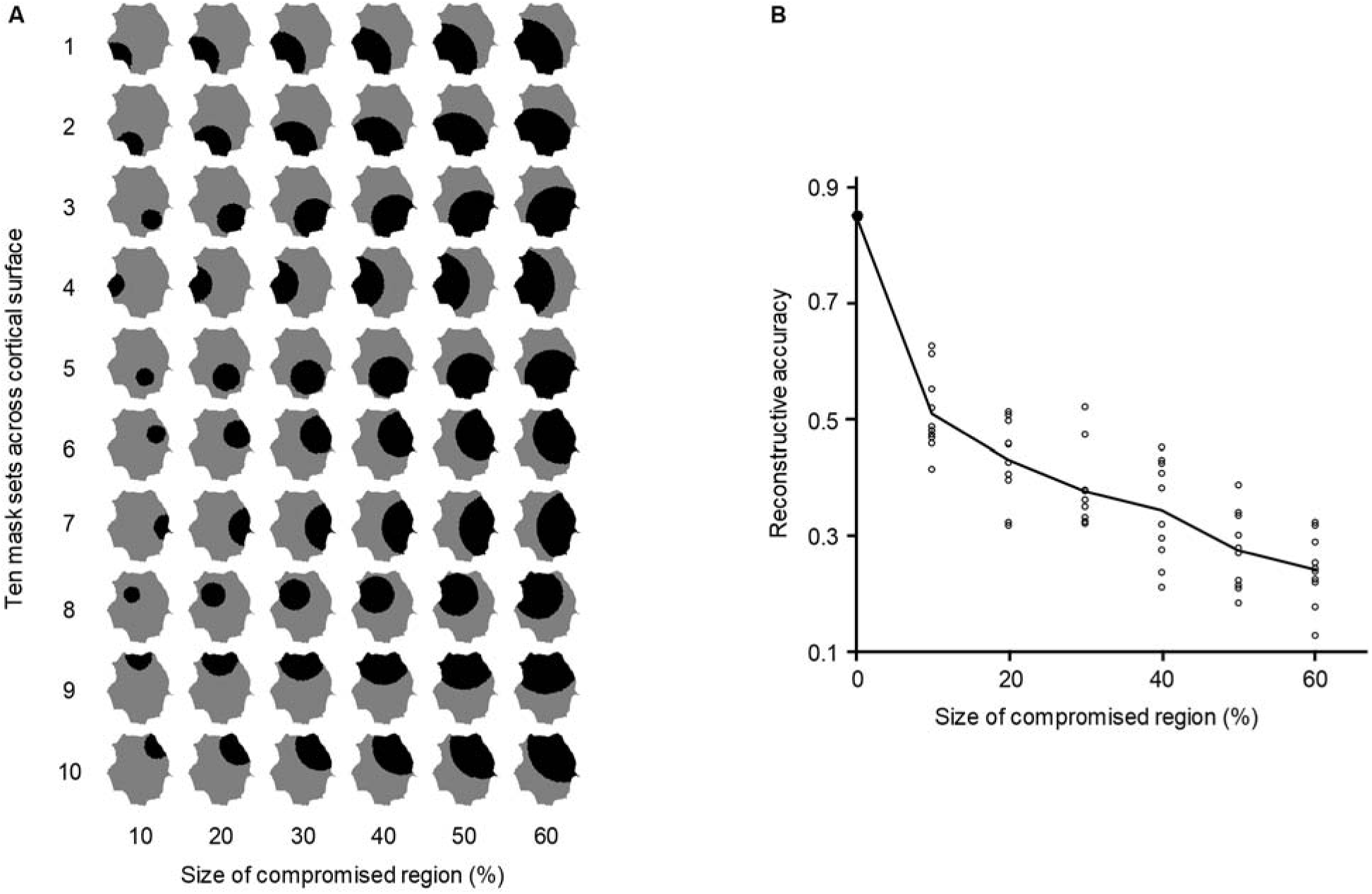
The reconstructive accuracy varies according to the size of the compromised area. **(A)** Ten vertices on the cortical surface were selected at random and served as the center of their respective compromised regions (indicated in black). From that center, we removed BOLD signals in incrementally larger regions, which spanned 10-60% of the cortical surface (with steps of 10%). The vertices nearest the centers were included in each of the masks. **(B)** This graph shows reconstructive accuracy as a function of the size of the compromised regions, as measured by calculating the average correlation between the original and reconstructed time series within that region. When we feed the DCGAN model an intact frame (size = 0%), the reconstructive accuracy is *r* = 0.85 ± 0.00. When the compromised region spans 10% of the cortical surface, the accuracy is *r* = 0.51 ± 0.07. From there, each incremental increase of 10% reduces the accuracy in a linear fashion (*F*(2.62,23.55) = 93.68, *p* < 0.001). Data points indicate the average reconstructive accuracy for each of the 10 cortical masks.

**Fig. S3.**
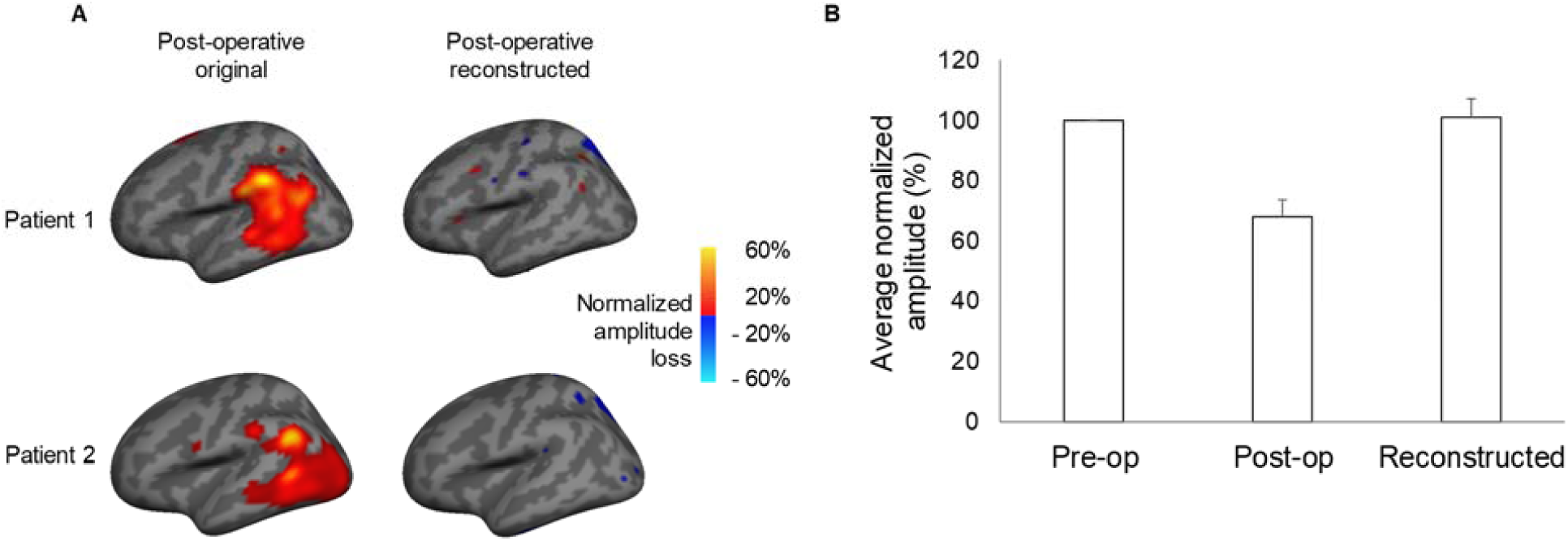
The BOLD signal amplitude is restored in the DCGAN-reconstructed frames. **(A)** We investigated whether there was any residual loss of BOLD signal in the DBS-compromised regions in the two patients with Parkinson’s disease after reconstruction. The left column shows the signal amplitude loss in the compromised region, calculated by subtracting the post-operative BOLD signal from the pre-operative BOLD signal. Both patients exhibit substantial signal loss following implantation. In contrast, the reconstructed BOLD signal shows no residual loss (right column). **(B)** The graph shows the normalized BOLD amplitudes within the compromised regions. The post-operative BOLD amplitudes are substantially and significantly lower than the pre-operative BOLD amplitudes (*t*(392) = −79.09, *p <* 0.001). The reconstructed BOLD amplitudes are significantly higher than the pre-operative BOLD amplitudes, but this increase is negligible (*t*(392) = 2.31, *p* = 0.02). Statistics were calculated based on data from both patients. The average signal amplitude increased from 67.84 (s.d. = 8.06) post-operatively to 101.06 (s.d. = 9.13) after reconstruction. Error bars represent standard errors.

**Fig. S4.**
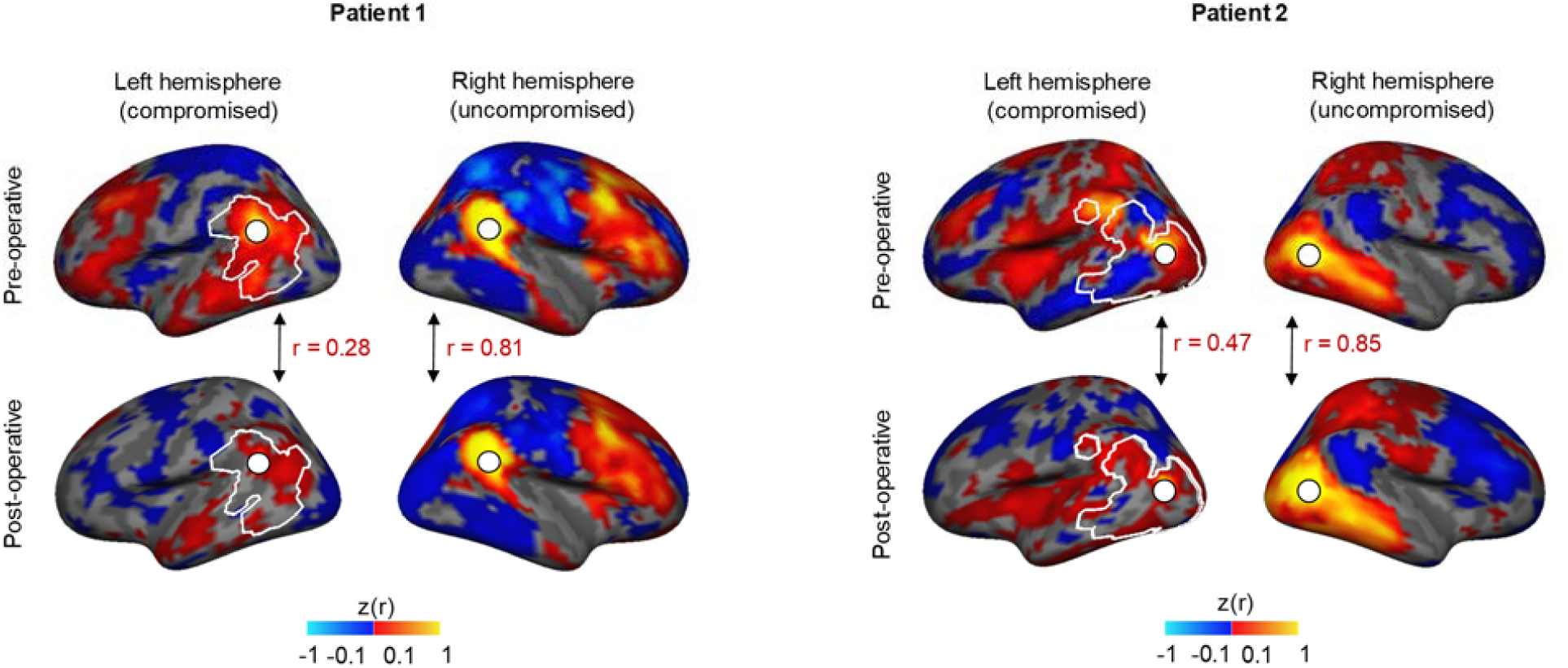
FC maps remain stable after electrode implantation. Here we show proof of concept that FC maps remain stable following electrode implantation surgery. We generated pre- and post-operative FC maps from one seed in the compromised region (left hemisphere), and the same seed in the uncompromised region in the right hemisphere. The left hemisphere post-operative FC map presents substantial differences from the pre-operative FC map, as indicated by a low correlation of *r* = 0.28 in Patient 1 and *r* = 0.47 in Patient 2. When looking at the uncompromised seed in the right hemisphere, the post-operative FC map is highly similar to the intact pre-operative FC map, as shown by a correlation of *r* = 0.81 in Patient 1 and *r* = 0.85 in Patient 2. Thus, functional connectivity does not seem substantially affected by electrode implantation.

